# Human lung explants as a predictive platform for evaluating the on-target, off-tumor toxicity of T cell bispecifics

**DOI:** 10.64898/2026.01.08.698374

**Authors:** Manuel Tschan, Tania Jetzer, Melanie Obenloch, Annika Blank, Carmen Yong, Nitya Nair, Lauriane Cabon, Elisa D’Arcangelo, Matthias P. Lutolf

## Abstract

T cell bispecifics (TCBs) are potent immunotherapies with proven efficacy in hematological cancers but limited success in solid tumors, where on-target, off-tumor toxicity has restricted their therapeutic index. Clinical discontinuation of TCB formulations against epithelial cell adhesion molecule (EpCAM TCB, Solitomab) and preclinical termination of a TCB against folate receptor alpha (FolR1-TCB) highlight the urgent need for predictive preclinical models to evaluate such toxicities. Conventional models, including cell lines and organoids, lack the immune and stromal complexity of the tumor microenvironment (TME), and while patient-derived explants preserve native tissue architecture and heterogeneity, quantification of TCB-mediated target cell killing has not been previously achieved in this system. Here, we establish a human lung explant model to assess TCB activity in matched tumor and normal adjacent tissues. Using a multi-modal approach combining flow cytometry, cytokine profiling, and multiplexed immunofluorescence, we show that EpCAM and FolR1 TCBs drive robust T cell activation and, critically, quantifiable epithelial cell killing above background levels. Normal lung explants exhibited consistently higher levels of killing than tumor counterparts, reflecting the on-target, off-tumor toxicities observed preclinically and clinically with the tested TCBs. In contrast, colon explants displayed poor viability *ex vivo*, limiting their suitability for assessing TCB-induced killing despite measurable immune activation. Our findings establish lung explants as a predictive and clinically relevant preclinical model that uniquely enables simultaneous quantification of TCB-mediated T cell activation and target cell killing. This model captures key features of TCB mechanism of action and recapitulates clinically observed toxicities, supporting its application for preclinical evaluation of TCB drug candidates and advancing the development of immunotherapies for solid tumors.

## Introduction

T cell bispecifics (TCBs) are a class of powerful immunotherapies that have shown striking results in hematological cancers^1–3^. However, for solid tumors, TCBs have faced a number of challenges, including on-target, off-tumor toxicity, which has resulted in significant adverse effects in preclinical and clinical models and has prevented achieving a suitable therapeutic index^4–6^. Notably, the clinical discontinuation of EpCAM TCB^7^ (Solitomab) and the termination of Folate Receptor 1 (FolR1) TCB during preclinical development, due to severe side effects in Non Humate Primates^8^, underscore the urgent need for robust preclinical models that can accurately predict and characterize off-tumor toxicity. Patient-derived explant cultures entail culture of fragmented donor tissue *ex vivo* and represent a compelling and clinically relevant model for preclinical cancer research^9–11^. Unlike conventional cell lines or organoid models, explants can maintain native tissue architecture and heterogeneity of cell populations within the tumor microenvironment (TME), including crucial immune cells^12–14^. This makes explants a highly patient-relevant, complex model for evaluating drug efficacy and toxicity ^9, 15^.

Indeed, explants have been successfully utilized for testing chemotherapies^16,17^, targeted therapies^18,19^, and immune checkpoint inhibitors^20^, though quantifiable metrics of target cell killing have often been lacking. Despite the proven capacity for T cell activation in explants^21–23^, the action of TCBs in explants has insufficiently been assessed^24^, nor has direct target cell killing in explants in the context of immuno-oncology compounds^12^.

To address this, we set out to investigate the activity of conventional EpCAM TCB, FolR1 TCB and CEACAM5 (Carcinoembryonic Antigen) TCB in human explant models of lung and colon tissues. Using a multi-modal approach in lung explants, comprising flow cytometry, secreted cytokine analysis and multiplexed immunofluorescence (mIF), we demonstrate that TCBs elicit a robust immune response and, importantly, that target cell killing is achieved in a quantifiable manner. Furthermore, we observe TCB on-target off - tumor toxicity reflected in lung explants: T cell activation takes place similarly in both tumor and adjacent normal explants, while epithelial cell killing takes place predominantly in tumor explants.

In contrast to lung, colon explants serve to highlight the limitations of the explant assay in tissues with low *ex vivo* viability. In these samples, target cell killing could not be reliably quantified using current explant protocols, underscoring the challenges of extending this approach to less viable tissue types.

## Results

### Characterization of lung explants

To investigate on-target, off-tumor effects of TCB treatment in explants, we first sought to understand the viability and high-level cell type composition in lung tissues sourced from multiple donors upon *ex vivo* culture. Characteristics of the donor patient population are summarized in SI Table 1. We focused on the following features of lung explant cultures: overall viability, maintenance of target expression levels *ex vivo* and major cell type composition (Fig. 1A).

**Figure 1.**
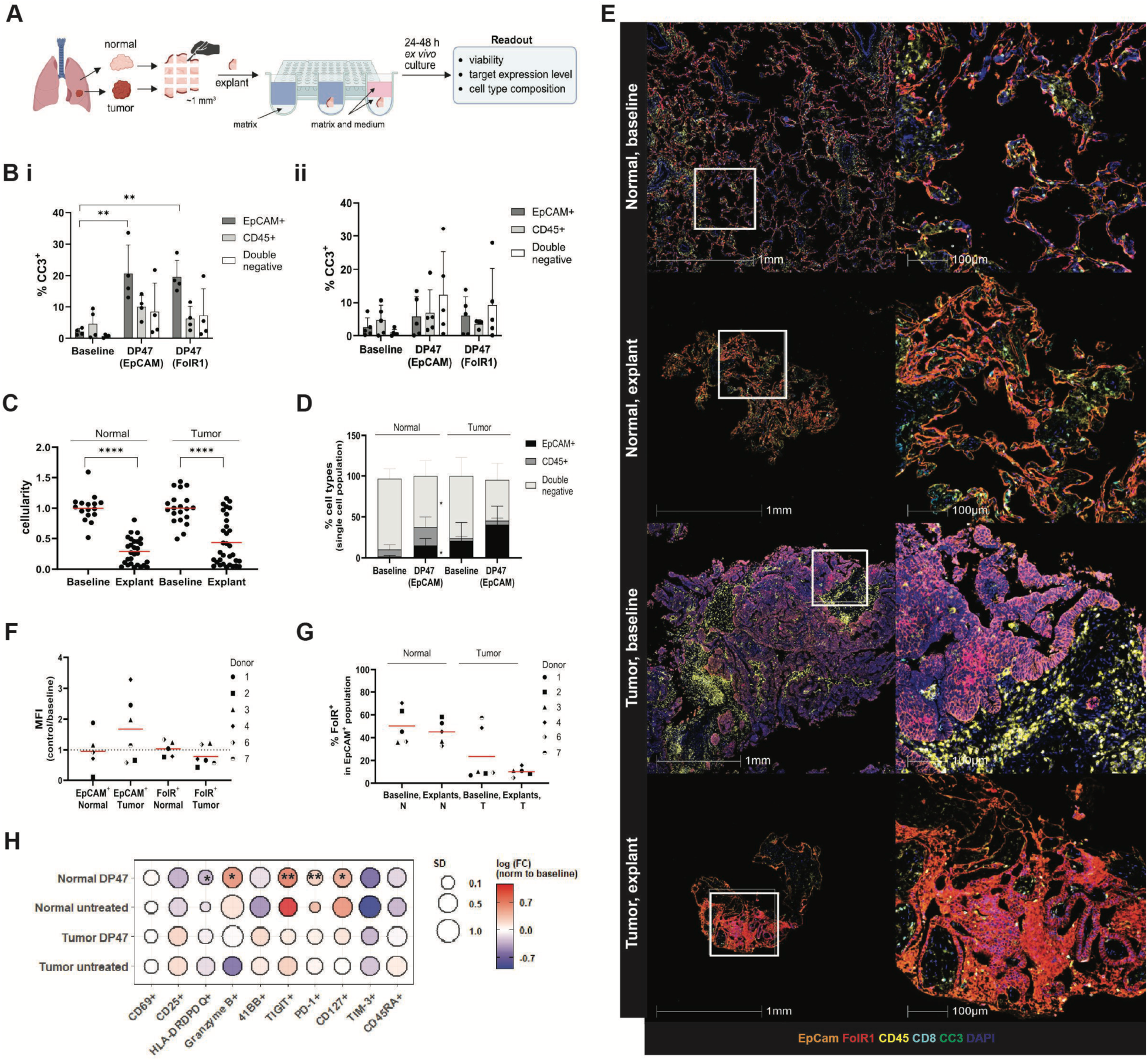
Characterization of lung explant viability, composition and antigen expression levels at 24-48 h ex vivo culture. (A) Schematic of workflow for lung explant generation, culture and readouts. (B) Apoptosis (% CC3-positive fraction) within EpCAM+, CD45+ and double negative cells in tissue digests from normal (Bi) and tumor (Bii) samples. Each dot represents an individual donor. (C) Cellularity measurements in baseline and matched explants, based on DAPI signal over tissue area in mIF images. Each dot represents one explant: min n= 4 explants/cohort/donor. (D) Gross composition in baseline and matched explants. (E) Representative mIF images of lung baseline and matched explants at 24h ex vivo culture. (F) EpCAM and FolR1 median fluorescence intensity (MFI) in control-treated explants compared to matched baseline tissues. (G) Overlap of EpCAM+ and FolR1+ expressing target cell populations in explants and baseline tissues. (H) Activation marker changes observed in untreated and control-treated lung explants compared to matched baseline tissues (n=3 donors for all groups except untreated normal lung explants, where n+ 2 donors). Red line indicates the distribution mean. Created in BioRender. Jetzer, T. (2026) https://BioRender.com/p3unogf.

A common drawback of explant models is their limited longevity *ex vivo*, with a typical reduction in viability within hours to days of culture^15,25,26^. Explant viability was assessed by quantifying apoptosis (cleaved-caspase 3, CC3) staining in epithelial (EpCam+), lymphocyte (CD45+) and double negative (EpCam-, CD45-) cell fractions at culture endpoint in control (DP47) -treated explants (Fig. 1B). Compared to baseline tissues, CC3 levels were higher in explants, in all sample fractions, for both tumor and adjacent normal tissues, with peak CC3 positivity observed in EpCAM+ cells in normal explants. Nevertheless, values didn’t exceed 20% across all groups on average.

We additionally checked whether CC3 positivity using flow cytometry correlated with tissue cellularity in fluorescently labelled sections of fixed explants. We quantified the number of nuclei present in baseline tissue and matched explants, which revealed a systematic, though highly variable, loss in cellularity of lung tissues upon 24 hr *ex vivo* culture, which exceeded apoptotic levels observed by flow (Fig. 1C). The discrepancy in explant viability assessed using these distinct approaches may be due to, on the one hand, the confounding effect of sample digestion in flow cytometry, which could lead to cell loss, and, on the other hand, potential dissimilarities between the amount of nuclei present within select histology slides, compared to the larger tissue amount used in flow cytometry, as well as the fact that the cell fraction positive for CC3 at a given timepoint does not reflect prior cellular losses. Nonetheless, tissue histology provides an additional compelling argument for the decreasing tissue viability in lung tumor and normal explants at 24 - 48 hours (h) *ex vivo* culture. Due to limiting amounts of patient tumor samples, especially for tumor tissue, comparisons were made between baseline and matched control-treated (DP47-treated) explants throughout this study, as untreated explants could only be generated for three donors. For these donors, by comparing DP47-treated to untreated explants, we observed that control treatment alone did lead to an increase in apoptosis levels within the epithelial fraction of normal explants, though the effect size varied drastically within the explant cohort (SI Fig. 1A). Characterized by a lack of tumor-targeting arm, but still containing a CD3-targeting arm, the DP47 control molecule may lead to antigen-independent T cell activation in this setting, resulting in elevated T cell cytotoxicity even in the absence of specific target binding; such effects have been previously described^27^. In terms of gross composition, all explants retained epithelial cells and lymphocytes throughout culture, while the double negative cell population showed a slightly decreasing trend (Fig. 1D). Similarly, in mIF stainings, lung explants consistently showed presence of epithelial cells, FolR1-expressing cells, CD45+ cells and cytotoxic T cells (CD8+) (Fig. 1E).

We next examined to what extent target expression (EpCam and FolR1) was retained in explants during culture using flow cytometry (Fig. 1F): compared to baseline tissues, lung explants from adjacent normal tissue-maintained target expression levels *ex vivo*; conversely, for explants obtained from tumor tissue, FolR1 levels on average dropped and EpCAM levels increased upon culture. Additionally, comparing normal to tumor explants, FolR1 levels were roughly comparable, while EpCAM levels varied more widely. The same assessment was done using an integrated density calculation, which takes into account both area and intensity of target expression in mIF images, showed a similar amount of signal variability, compared to baseline tissues, as flow cytometry (SI Fig. 1B), strengthening the notion that target expression levels are in aggregate similar to baseline levels (though with high variability between donors). With the aim of applying both EpCAM TCB, as well as FolR1 TCB to lung explants, we next asked whether all putative target cells indeed co-expressed both proteins. This data can inform the amount of intended target cells present (i.e. tumor cells expressing EpCAM and/or FolR1) vs number of cells expressing the target molecule, but which are not epithelial cells (Fig. 1G). The overlap of EpCAM+ and FolR1+ cells in normal explants was high (roughly 50% for both), as expected; interestingly, this was not the case for tumor explants, where FolR1 levels in the EpCAM+ population were rather low (roughly 25%), indicating a heterogeneous expression on tumor cells leading to downregulation or survival benefits in antigen-negative fractions during culture or from the outset. Collectively, these data indicate that in explants generated from matched lung normal and tumor tissues, viability declines from baseline to assay endpoint of 24-48h, likely limiting the usability of lung tissue for drug assays requiring substantially longer culture times *ex vivo*. Gross sample composition remained stable over time, with key cell populations being present within explants, including target-expressing epithelium, CD45 + cells and CD8 + lymphocytes.

We then determined whether *ex vivo* culture could modify T cell activation status. Granzyme B (GrzB) positivity, as well as expression of TIGIT, PD-1 (programmed death 1) and CD127 were moderately increased in CD4+ and CD8+ cells within normal lung explants, compared to baseline tissues, indicating changes in T cell activation and exhaustion patterns because of *ex vivo* culture. This trend was further increased upon treatment with the DP47 control. For tumor explants, notable culture- or DP47-induced changes in T cell activation included CD25, GrzB, TIGIT and 4-1BB (Fig. 1H). Further, HLA-DRDPDQ and TIM-3 were downregulated in both tumor and normal explants, compared to their baseline tissues, while CD69 remained unchanged in both explant groups. These data suggest that *ex vivo* culture itself induced changes in T cells, potentially comprising low level activation, increased effector function and partial exhaustion.

### TCB treatment leads to immune cell activation in lung tumor and normal explants

We next set out to examine the effect of conventional TCB treatment on immune cell activation in matched lung tumor and normal explants, whereby treated explants were compared to each other, as well as to baseline tissues (Fig. 2A). Importantly, application of EpCAM or FolR1 TCBs in lung explants (and CEACAM5 TCB in colon explants, see Fig. 5) did not systematically alter the frequency of EpCAM+, FoLR1+ or CEACAM5+ cells detected by flow cytometry, respectively.

**Figure 2.**
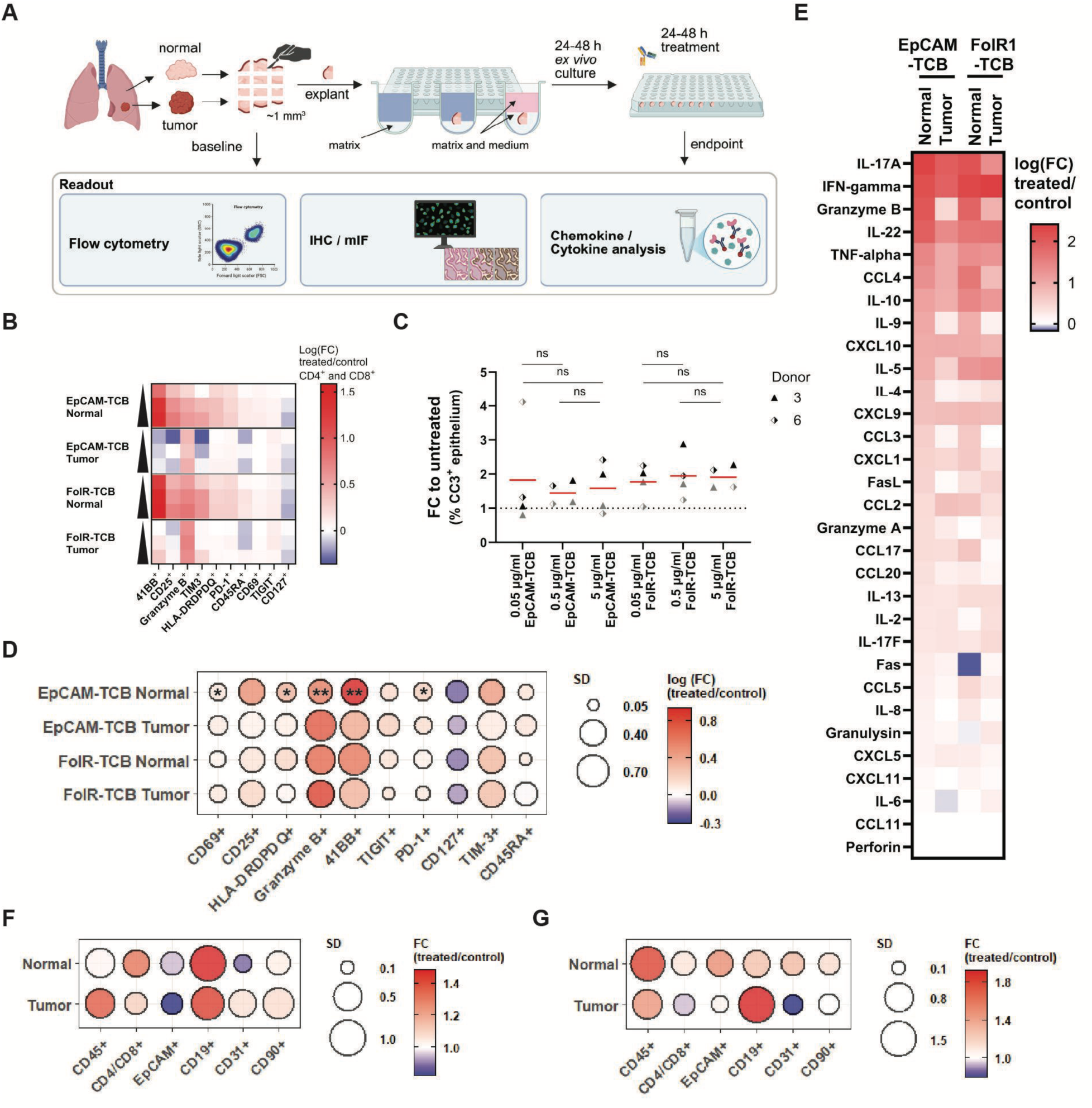
TCB treatment leads to immune cell activation in lung explants. (A) Schematic of workflow for *ex vivo* treatment and readouts comparing baseline tissues and matched explants. (B) TCB dose-dependent changes in explant immune activation markers in CD4+ and CD8+ T cells. (C) Epithelial cell apoptosis in lung normal (grey) and tumor (black) explants. (D) Changes in immune activation markers and (E) secreted factors in lung normal and tumor explants, compared to controls, upon TCB treatment. (F) Changes in population fractions in lung normal and tumor explants, compared to controls, upon EpCAM TCB treatment and (G) FolR1 TCB treatment. Red lines indicate distribution mean. Created in BioRender. Jetzer, T. (2026) https://BioRender.com/p3unogf.

To determine an appropriate dose for our TCB treatments, we first tested the effect of a range of doses for both EpCAM and FolR1 TCBs on T cell activation and CC3 positivity compared to control. Both tumor and normal explants showed only minor (up to a fold change of 1.6) changes in T cell activation markers, in both normal and tumor explants at increasing TCB doses (Fig. 2B). However, killing (as quantified by CC3 positivity within the epithelium) remained statistically unchanged, especially at higher doses tested (Fig. 2C), indicating that a dose of 0.5 ug/ml was sufficient to observe the maximum achievable killing effect in this system for both TCBs. Changes in T cell marker expression trended similarly between both TCB treatments: normal lung explants were characterized by pronounced upregulation of CD25, HLA-DRDPDQ, GrzB, 4-1BB, TIGIT, TIM-3 and PD-1, while downregulating CD127, altogether indicating robust T cell activation and a concomitant shift towards terminal differentiation and exhaustion. Despite inter-donor variability (SI Fig. 2A), we observed a general trend in tumor tissues mirroring what we observed in normal tissue, except for higher GrzB levels (Fig. 2D). Marker levels in baseline tissues are summarized in SI Fig. 2B; of note were TIGIT levels at baseline, which were strongly elevated in the tumor cohort, suggesting that tumor tissues contained a higher fraction of exhausted T cells, compared to normal tissues, a feature typical of lung cancer (ref).

Analysis of secreted factors in treated vs control-treated explant medium further corroborated these observations (Fig. 2E). The top three factors secreted in treated normal explants (for both TCB treatments) were pro-inflammatory cytokines IL-17A, IFN-𝜸 and GrzB, indicating a potent cytotoxic T cell response. Notably, increased IFN-𝜸 serum levels were observed clinically 24-48 h upon treatment with the EpCAM TCB Solitomab^7^. For treated tumor explants, IL-17A, IFN-𝜸 and IL-22 were highest, though GrzB was still elevated. The secretion of additional pro-inflammatory, regulatory, and chemokine molecules were also increased, such as TNF-α, CCL4, IL-10, and cytokines involved in IFN-𝜸 signaling (CXCL10 and CXCL9). Of note was the paucity of perforin, which may reflect the added presence of a non-cytolytic signaling function of granzymes in this system^28,29^, as well as perforin-independent target cell killing, as seen in other contexts^30–32^. The lack of a complete immune response, due to inability to recruit additional immune cells, could further explain the low abundance of the broad pro-inflammatory cytokine IL-6, as well as other chemokines. Finally, we profiled the effect of TCB treatments on other cell populations present within explants (Fig. 2F, G). In all conditions examined, only FolR1 TCB treatment in tumor explants led to a meaningful population increase (fold change > 1.5) in CD19+ cells, though with high variability, and a decrease in CD31+ cells. An enrichment of B cells has been reported in lung cancers before^33,34^; while CD19+ cells in this model may likely represent tissue-resident B cells, which plausibly could expand due to the local activated cytokine environment or interaction with T cells following TCB treatment^35^. Collectively, our analysis of both secreted factors and T cell markers in lung explants confirmed that TCBs elicited strong T cell activation in both tumor and normal tissues. Furthermore, normal explants were on par with tumor explants in terms of T cell activation, in line with on-target but off-tumor activity previously reported with these drugs.

### Normal explants exhibit higher TCB-induced killing than tumor counterparts

Explants generated from a range of tissues have thus far been utilized to assess the effects of immuno-modulatory compounds such as checkpoint inhibitors *ex vivo*^20^ and are indeed considered a robust assay for characterizing T cell activation. However, studies quantifying killing/death of epithelial target cells in explants treated with immunomodulatory compounds are scarce. This paucity of killing data is likely due to poor explant viability at assay endpoint, which confounds killing readouts. We therefore examined whether quantitative killing could be measured in our model setup, which is characterized by a short assay duration, reasonable retention of target levels and overall viability at endpoint. Positivity for CC3 was detected using flow cytometry and was higher in EpCAM+ cells in EpCAM TCB-treated normal explants, compared to matched DP47-treated controls (Fig. 3A). These levels substantially exceeded the levels of cell death induced by *ex vivo* culture alone, confirming the TCB-mediated killing effect. Critically, CC3 positivity in normal epithelium was not only high by comparison to the tumor explant cohort; rather, but it was also high in absolute terms: for TCB treatments, approximately 50% of epithelial cells in normal explants were undergoing apoptosis at endpoint (Fig. 3A).

**Figure 3.**
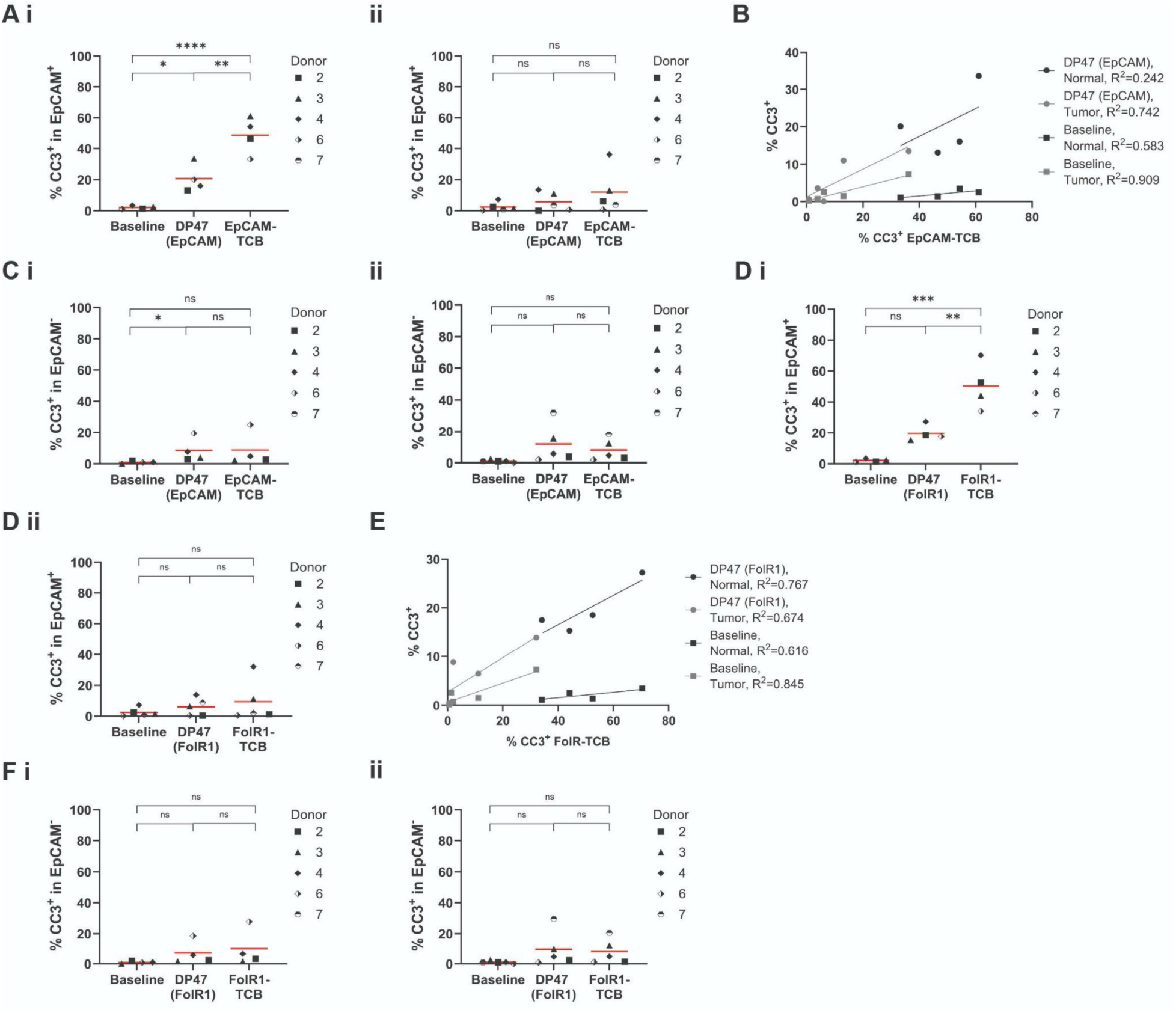
TCB-induced target cell killing is detected in lung explants by flow cytometry. (A) Percentage of CC3+, EpCAM+ cells upon EpCAM TCB treatment in matched baseline tissues, control-treated and TCB-treated explants, for normal (I) and tumor (II) samples. (B) Correlation between CC3 positivity in control-treated explants, EpCAM TCB-treated explants and matched baseline tissues. (C) Percentage of CC3+, EpCAM- cells upon EpCAM TCB-treatment in matched baseline tissues, control-treated and TCB-treated explants, for normal (I) and tumor (II) samples. (D) Percentage of CC3+, EpCAM+ cells upon FolR1 TCB treatment in matched baseline tissues, control-treated and TCB-treated explants, for normal (I) and tumor (II) samples. (E) Correlation between CC3 positivity in control-treated, FolR1 TCB-treated explants, explants and matched baseline tissues. (F) Percentage of CC3+, EpCAM- cells upon FolR1 TCB treatment in matched baseline tissues, control-treated and TCB-treated explants, for normal (I) and tumor (II) samples. Red lines indicate the distribution mean.

Focusing on EpCAM TCB, we next asked whether CC3 positivity in baseline or control-treated samples could predict the magnitude of killing observed in treated samples and found that in tumor, both metrics roughly predicted the killing effect upon TCB treatment (while for normal tissues only CC3 levels in control-treated explants were predictive) (Fig 3B). This suggests that the CC3 positivity at baseline, as well as in control-treated explants, reports on an immunologically active state present in the absence of treatment, or CD3 engagement occurring upon DP47 treatment. The finding that *ex vivo* culture itself induces expression of numerous T cell activation markers in lung explants above baseline corroborates this hypothesis (Fig. 1H).

Within the EpCAM-negative cell fraction in EpCAM TCB treated explants we did not observe increased CC3 levels beyond those observed in DP47 controls, further indicating that the treatment- induced killing was specific to the target epithelial population (Fig. 3C). Strikingly however, though T cell activation metrics were similar in tumor and normal explants (Fig 2E, F), TCB-induced killing was significantly lower in tumor explants, compared to normal counterparts, specifically within the EpCAM+ cell population (Fig 3A). This may reflect the excluded/deserted nature of tumor baseline tissues (SI Table 1), the increased exhaustion of tumor T cells at baseline (as shown by elevated TIGIT positivity), as well as other potential tumor intrinsic TCB-resistance mechanisms ^36^. In line with this, lower CC3 levels were already present in control-treated tumor explants, compared to control-treated normal explants (Fig. 2B). Near-identical trends were observed for FolR1 TCB- treated explants (Fig. 3D-F), whereby normal explants again showed improved killing, compared to tumor explants, and that CC3 positivity in baseline and control-treated tissues could inform on the extent of killing achieved upon TCB treatment. Additionally, FolR1 expression was found to be lower in tumor explants, compared to normal explants, which may also have contributed to lower target killing in response to FolR1 TCB in the former (Fig. 1G).

Overall, by assessing CC3 as killing readout in lung explants, we established that TCB treatment leads to a robust increase in target cell killing in normal explants, while killing in tumor tissues was likely hampered by a combination of paucity of T cells within epithelial areas, antigen expression levels and other, tumor-intrinsic resistance mechanisms.

### TCB-induced killing displays spatial and donor variability in lung explants

T cell mediated target cell killing in solid tissues is governed by local TME features, including neighboring cells and stromal architecture^37–39^. We next explored the possibility of observing such killing heterogeneity in our setup, both between donors, but also across explants from the same donor. Using mIF, we observed epithelial cells, T cells and CC3 signals within the spatial tissue context (Fig 4A). Aggregated CC3 fluorescence values showed an increasing trend in treated explants from normal tissue (Fig. 4Bi), while there was no significant difference for tumor explants (Fig. 4Bii). Similar results were also observed in FolR1 TCB treated explants (Fig. 4C). Specifically, apoptosis showed a ∼ 3-4-fold change difference between control and treated normal explants. These data were therefore consistent with the analysis of CC3 positivity by flow cytometry (Fig 3A, D). A clear advantage of examining fluorescent images of individual explants over flow cytometry, however, was the possibility of gaining insight into the spatial heterogeneity of the TCB-induced killing response across fragments. Indeed, when examining CC3 positivity across donors this way, we found a striking degree of intra-patient heterogeneity with respect to apoptosis in normal explant cohorts (SI Fig. 3A, B showing donor 2 and 4, respectively), while explants generated from tumor tissue showed lower CC3 signal and signal heterogeneity in each FFPE slide.

**Figure 4.**
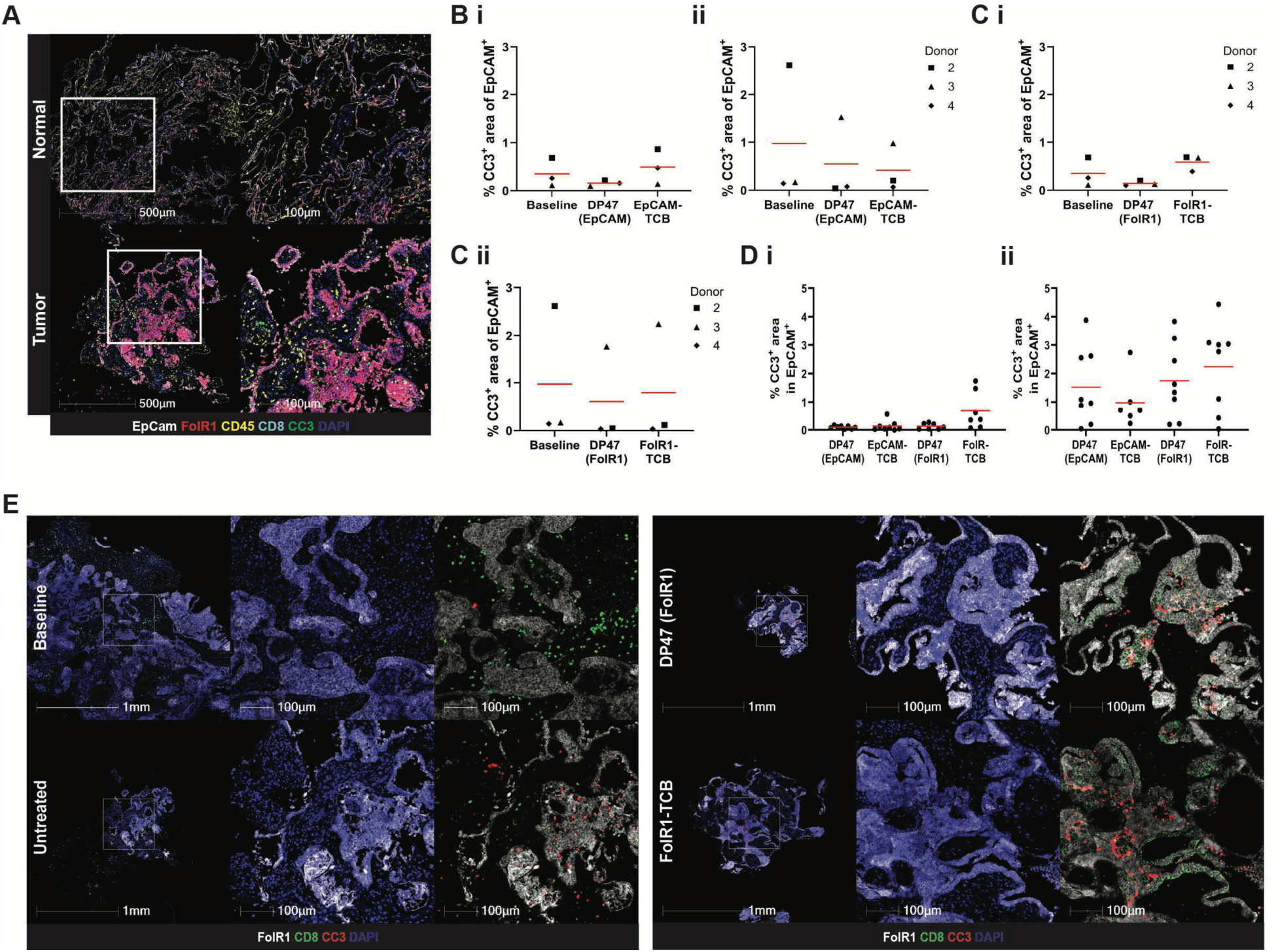
TCB-induced target cell killing is detected in lung explants by immunofluorescence. (A) Representative image of control-treated and EpCAM TCB-treated lung normal explants showing target-expressing cells, CD45 positive, T cells and CC3 staining. (B) Inter-donor variability of aggregated CC3-positive epithelial area in baseline, control-treated and EpCAM TCB-treated lung explants. (C) Inter-donor variability of aggregated CC3-positive epithelial area in baseline, control-treated and FolR1 TCB-treated lung explants. (D) Intra-donor variability of CC3-positive epithelial area in baseline, control-treated and TCB-treated explants from donor 3. (E) Representative images of donor 3 tumor tissue and explants showing FolR1-expressing cells, CD8 T cells and CC3 staining. Red lines indicate the distribution mean.

**Figure 5.**
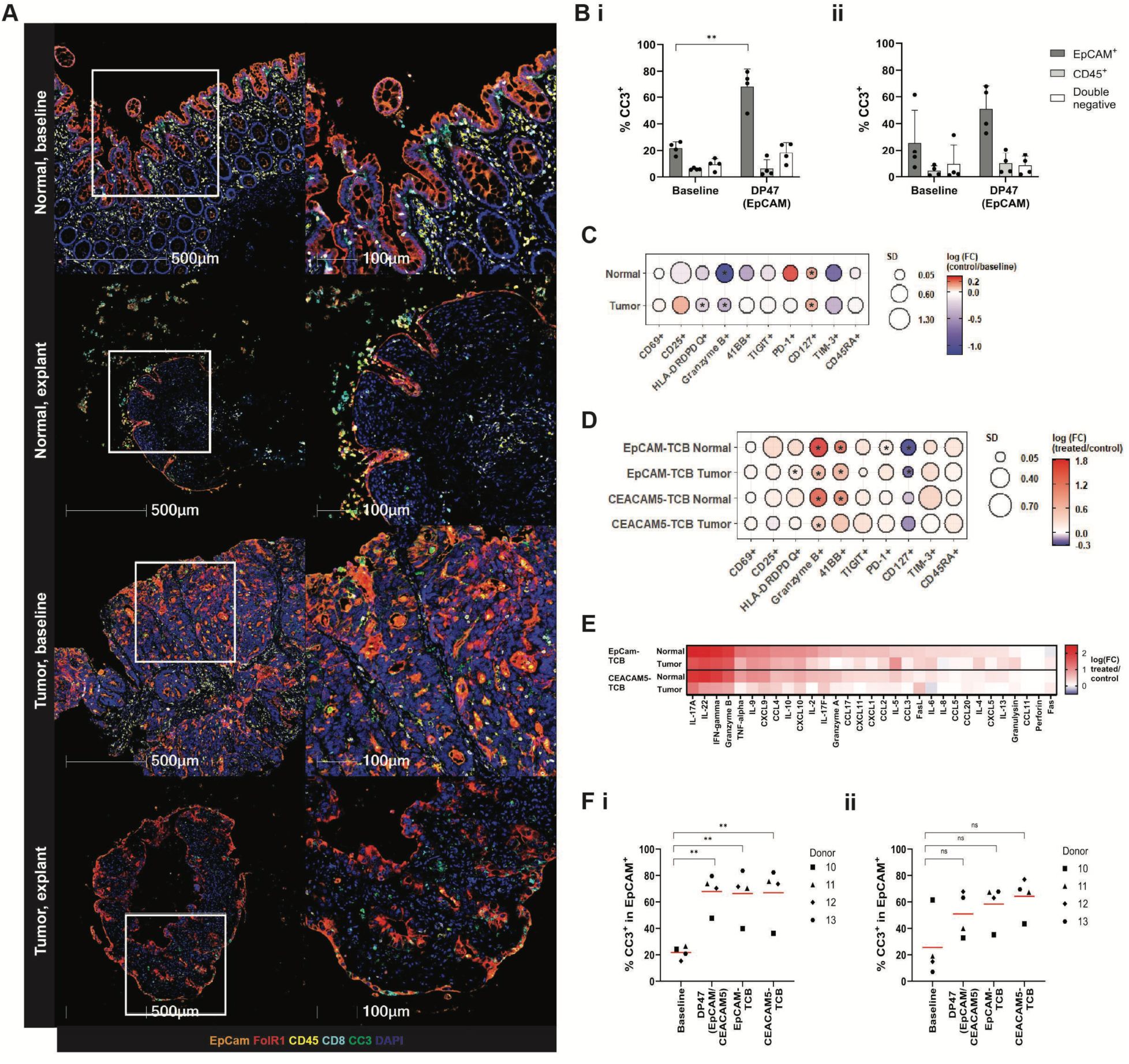
TCB treatment induces Immune activation but no detectable target cell killing in colon explants. (A) Representative mIF images of colon baseline and matched explants at 24h *ex vivo* culture. (B) Apoptosis in baseline colon tissues and matched explants as %CC3+ fraction within EpCAM+, CD45+ and double negative fractions of tissue digests. (C) Changes in activation markers observed in colon explants compared to matched baseline tissues, due to *ex vivo* culture. (D) Changes in immune activation markers in colon normal and tumor explants, compared to controls, upon TCB treatment. (E) Changes in chemokines/cytokines in colon normal and tumor explants, compared to controls, upon TCB treatment. (F) Inter-donor variability of aggregated CC3-positive epithelial area in control-, EpCAM TCB- and CEACAM5 TCB-treated colon explants. Red lines indicate the distribution mean.

For donor 3, and especially in the FolR1-treated cohort, CC3 signal was elevated robustly in tumor explants, compared to control (Fig. 4D); this was not obvious in our assessment of CC3 positivity by flow cytometry. As donor 3 had been categorized as immune excluded at baseline (SI Table 1), we were intrigued as to whether this donor displayed a more infiltrated phenotype at culture endpoint in the FolR1 TCB-treated cohort (Fig 4E). We made two notable findings for this donor: firstly, CD8+ T cells, which were present exclusively in stromal areas in tumor baseline tissue, were localized within epithelial zones upon FolR1 TCB treatment and additionally appeared more abundant and prominently integrated throughout the epithelium. These observations suggest that for this donor, T cell infiltration into tumor epithelium upon activation was indeed possible and led to subsequent target cell killing, thereby exemplifying the desired mode of action within tumor tissue (the only such case in our tumor cohort). Consistent with this, among donors 2- 4 analyzed, donor 3 showed the lowest amount of TIGIT and TIM-3 exhaustion markers in tumor explants upon TCB treatment (SI Fig. 2B). Importantly, explants from this donor did also not suffer from other culture-induced caveats such as loss in overall or epithelial cellularity. Secondly, CD8+ T cell recruitment into tumor epithelium could also be observed in DP47-treated tumor explants for donor 3, though to a lesser extent than in the FolR1 TCB-treated condition (Fig. 4E), suggesting, surprisingly, that DP47 stimulation (and therefore activation of T cells via CD3) could lead to T cell infiltration into epithelial areas.

By contrast, an examination of mIF images from donor 2 and 4, showed a lack of CC3 positivity in tumor explants (consistent with mIF image quantification (Fig. 4C ii), as well as flow data for these two donors (Fig. 3C ii)). Specifically, tumor fragments, as well as baseline tumor tissue, from donor 2, showed a lack of FolR1 antigen expression (SI Fig. 3C), which had been observed to be low in flow cytometry for this cohort (Fig. 1F). In contrast, donor 4 presented higher antigen levels, but also a paucity of CD8+ T-cells (SI Fig. 3D). This mIF analysis of donors 2 and 4 strengthened hypotheses on the reasons behind lack of tumor cells killing for these donors, namely low antigen expression and insufficient effector T-cells, respectively.

While the mIF approach was far more subject to inter-fragment variability compared to flow cytometry, an equivalent level of statistical significance could likely be achieved with additional staining on the same explant FFPE blocks (i.e. by considering a more representative volume for each explant). Importantly however, this analysis allowed us to perform a deeper characterization of specific fragment cohorts of interest, such as those from donor 3, thereby revealing spatial heterogeneity in treatment response and visualizing T cell infiltration into epithelium upon treatment.

### TCB treatment induces immune activation but no detectable target cell killing in colon explants

The characterization of explants from lung normal and tumor tissues upon TCB treatment was enabled by the combination of favorably short drug action times and relative longevity of lung tissue fragments *ex vivo* (i.e. less than 30% CC3+ epithelial cells at assay endpoint as assessed by flow cytometry, Fig. 1B).

To ascertain whether the same readouts could be collected from tissue types with poorer *ex vivo* viability, we applied TCB treatments to matched colon tumor and normal samples (donor details are listed in SI Table 1). Of note, two doses (0.5 ug/ml and 5 ug/ml) for CEACAM5 TCB and matching DP47 control were tested between three colon tissue donors and data were pooled across these after observing no differences in T cell activation or killing. CEACAM5 is highly expressed in colonic epithelial cells and colon cancer and was therefore chosen in this study, along with EpCAM TCB. For colon explants, *ex vivo* viability at endpoint (24 or 48h) was much lower than in lung explants, with tissue integrity being evidently impacted by *ex vivo* culture (Fig. 5A) and CC3 positivity reaching 80% in epithelial cells (Fig. 5B). DP47 treatment itself was again observed to influence immune activation markers in colon, though differed from lung in terms of the specific markers impacted: PD-1, CD127, GrzB and TIM-3 were most differentially expressed (Fig. 5C). Most notable was the downregulation of GrzB in control-treated colon explants, which may be directly influenced by the state of “tolerance” typical for T cells within the colonic lamina propria *in vivo*^40^. Conversely, upon treatment with EpCAM or CEACAM5 TCB, GrzB and 4-1BB were the top 2 activation markers enriched in both tumor and normal colon explants as assessed by flow cytometry, followed by CD25 and TIM3 (Fig. 5D). Much higher levels of GrzB and 4-1BB were observed in normal explants, where expression fold changes reached 48 and 19 for GrzB and 4-1BB, respectively; in contrast, in tumor explants fold changes amounted to 3.6 and 5.7. Overall and similarly to lung explants, this pattern confirmed that T cells are activated in colon explants upon TCB treatment. This conclusion is further corroborated by the finding that IL-17A, IL-22, GrzB and IFN-𝜸 were secreted in both normal and tumor explants upon TCB treatment (Fig. 5E), with the overall chemo-and cytokine secretory profile in colon explants bearing strong similarities to lung explants.

Finally, we examined whether treatment-induced killing could also be quantified in colon explants. For this, we assessed CC3 positivity in EpCAM+ cells using flow cytometry at endpoint. No significant differences were observed between control- and TCB-treated normal or tumor colon fragments, though a non-significant trend (NS; p=x) was present in the latter (Fig. 5F). Importantly, control-treated explants already showed strongly elevated levels of apoptosis, likely reflecting the high levels of culture-induced damage, thereby making it a challenge to observe treatment-induced add-on effects in target cell death.

These data exemplify the usability of explants as an immune activation assay despite severely deteriorated tissue architecture and (especially epithelial) viability in specific tissue types. In these same tissues however, explants are poorly suited to capture drug-induced killing and numerous studies utilizing explants from such tissue types fall short in capturing this functional readout.

## Discussion

We here establish a robust *ex vivo* lung model for evaluating the on-target, off-tumor toxicity of conventional EpCAM and FolR1 TCBs. By combining a short assay duration with multi-modal analysis, we show that this model successfully captures and quantifies TCB-induced immune responses and cytotoxic effects within the context of a complex, all-human and patient-specific model of solid tumor and normal adjacent tissue. This work exemplifies how lung explants can be employed to support rapid risk assessment of drug candidates during preclinical development.

Our findings confirm that lung explants, despite a modest loss in cellularity *ex vivo*, maintain architecture and key cell populations, including target-expressing epithelial cells and immune cells. While the observed decline in viability represents a limitation for long-term studies, it does not, in the case of lung tissue, impede short-term assays. Lung explants from both normal and tumor tissues are similarly immunologically active, with TCBs inducing a robust T cell activation profile in both. This response, characterized by the upregulation of activation markers like GrzB and 4-1BB and a concurrent increase in exhaustion markers such as PD-1 and TIM-3, significantly exceeded any changes in immune cell activation stemming from the *ex vivo* culture and DP47 treatment. Interestingly, the lack of other cytotoxic proteins like perforin and granulysin suggested a perforin-independent killing pathway was engaged upon TCB treatment, which is known to rely on receptor endocytosis^30–32^. The secretion of pro-inflammatory cytokines and chemokines in explant supernatants further corroborates the potent TCB-mediated T cell engagement.

A central finding of this study is a meaningful quantification of target cell killing using flow cytometry and mIF, a major challenge in previous explant research, where killing was simply inferred, based on T cell activation^13,14,21,23^. Critically, while immune activation data showed similar trends between normal and tumor explants, killing data revealed much higher levels of treatment-induced, T cell mediated apoptosis in lung normal explants, compared to tumor counterparts. This was in line with the assessment of tumor immune infiltration status, which qualified tumor tissues included in this study as immune deserted or excluded, as well as other potential immunosuppressive mechanisms within explant TMEs. The observation of substantial target cell killing in fragments from adjacent normal tissue constitutes the first report of on-target off-tumor T cell mediated killing quantified in explants. Consequently, this assay presents a valuable opportunity to screen TCB drug candidates for greater tumor specificity while minimizing normal tissue reactivity, potentially leading to an enhanced therapeutic index in clinical settings.

It is estimated that approx. 10-30% NSCLC tumors are immune deserts while 40-60% are immune excluded^41^. We believe it to be likely that T cells in our tumor explant cohort became activated within the stroma of tumor explants both through minimal contact with target-expressing cells, as well as through the saturating levels of TCB used. Indeed, we observed that killing plateaus at higher TCB doses. Moreover, this activation did not translate into effective target killing in tumor explants, highlighting a critical distinction between T cell activation and T cell effector function^42^ within a complex, solid 3D tissue microenvironment. In our sample cohort, it remains unclear whether this occurred, due to poor accessibility or additional TCB resistance mechanisms^36^; however, these killing data validate the potential to use our lung explant model to capture TME features with high relevance for drug efficacy. Additionally, our data compellingly shows the on-target, off-tumor toxicity of both EpCAM and FolR1 TCBs, which were both able to elicit robust killing in normal lung tissue. Likely, the killing of normal epithelium in this model results from antigen levels and T cell states conducive to successful TCB function, coupled with the absence of TCB resistance mechanisms.

Beyond aggregate quantification of CC3 signal, the mIF data provides a powerful spatial dimension to our findings. Specifically, it confirms the trends from flow cytometry while also underscoring a high degree of intra-patient heterogeneity between fragments from the same tissue and donor. While it is possible that differences in *ex vivo* viability between fragments underlie this finding, we believe it also to be likely that such heterogeneity also reflects regional differences in tissue properties such as target cell abundance, antigen expression and T cell infiltration and fitness at the time of fragmentation. We therefore examined tumor explants from donor 3 in more detail, which revealed that, despite being categorized as immune excluded at baseline, FolR1 TCB (and to a weaker extent DP47 treatment) led to CD8 T cell recruitment into tumor epithelium and increased target cell killing, showcasing successful re-direction and engagement of T cells for this tumor and the value of our multi-modal approach for investigating killing in this model. Further analyses comprising additional explant FFPE slices from these samples could improve the strength of the conclusions drawn from our mIF data.

Finally, we applied our explant workflow to colon explants, which showcased the model’s limitations. Colon tissue appeared to suffer much more considerably from declining viability *ex vivo*, compared to lung, within the epithelial cell fraction specifically (while viability in immune cells remained high at endpoint). Thus, while T cells in explants from colon normal and tumor could be activated by conventional EpCAM and CEACAM5 TCBs (suggesting again the possibility of on-target, off-tumor cytotoxicity in this model), the high background apoptosis levels made colon an unsuitable system for quantifying killing per se. The applicability of explant protocols across tissue types therefore highly depends on the choice of readout and how this is impacted by the intrinsic viability and longevity of the tissue type, a key consideration for future studies.

Overall, we established the lung explant model as a translatable preclinical tool for assessing TCB activity in a patient-true, all-human system, with capabilities to quantify immune cell activation and target cell killing. We have shown that the model recapitulates the on-target, off-tumor toxicity associated with TCBs in solid tumors.

This study suffers from several limitations, most of which are inherent to the fact that explants recapitulate the compositional variability observed both within the patient cohort chosen, and within any given tissue, therefore making it challenging to attain statistical significance in quantitative assays. To counteract this, a higher number of donors could be included in future studies. Including donors with well-infiltrated tumor tissues will allow a characterization of killing levels attainable in such tumors, as well as assessment of potential TCB-resistance mechanisms other than poor immune infiltration. The lung normal tissue utilized in this study was obtained from lung areas at variable distance from tumors and therefore constitute normal adjacent tissue. We believe that an examination of true normal tissue is warranted (for example from cadaveric donors); similarly exploring on- target off-tumor effects in secondary organs (also potentially characterized by high target expression) would represent a valuable contribution from a safety assay standpoint. Next, although explants from lung maintained relatively high viability levels, a major limitation of the explant field is also reflected in this work: testing of longer drug regimens, combination treatments, repeated dosing or assessments of the development of adaptive resistance mechanisms are currently unfeasible strategies across tissue types, due to insufficient explant longevity. Lastly, explants in their current culture format cannot recapitulate systemic immune responses, including cytokine release syndrome, and biodistribution of TCBs, all of which are important factors influencing TCB efficacy and safety profiles.

Efforts to prevent or curb tissue viability loss *ex vivo* (across tissue types, but especially for tissues such as colon) are very much required, to capitalize on the predictive power of explant tissues for the assays described. Specifically, modulation of culture parameters such as medium composition and oxygenation remain areas of intense work. In the short term, such studies will clarify the signaling underlying poor explant *ex vivo* fitness; in the long term, explant models that can sustain longer *ex vivo* culture may be used for longer or combinatorial drugging regiments while still offering an unmatched capacity to model tissue-level therapeutic responses.

**Supplementary Figure 1.**
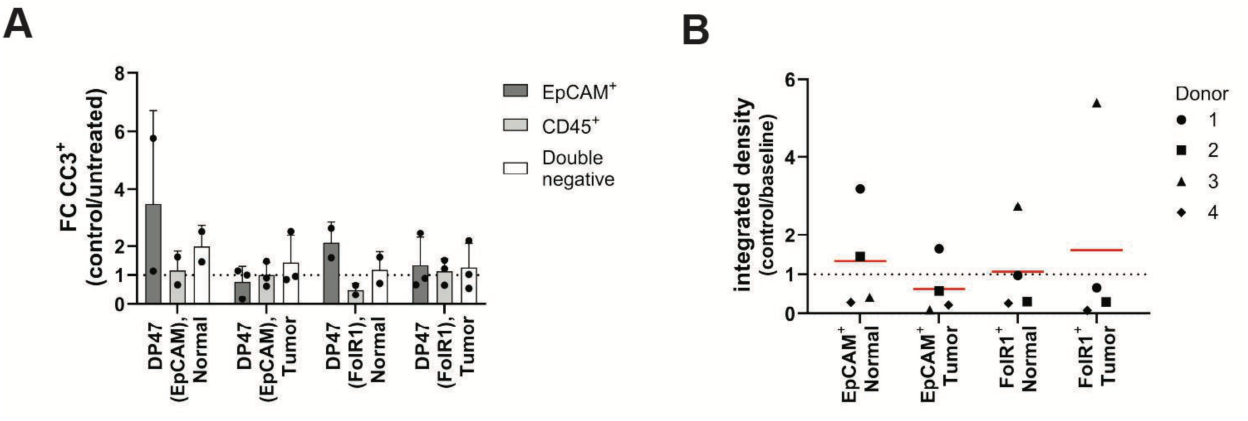
(A) Fold change (FC) comparison of lung explant viability between untreated and control-treated explants (B) EpCAM and FolR1 expression in mIF images of fixed explants, compared to matched baseline tissues.

**Supplementary Figure 2.**
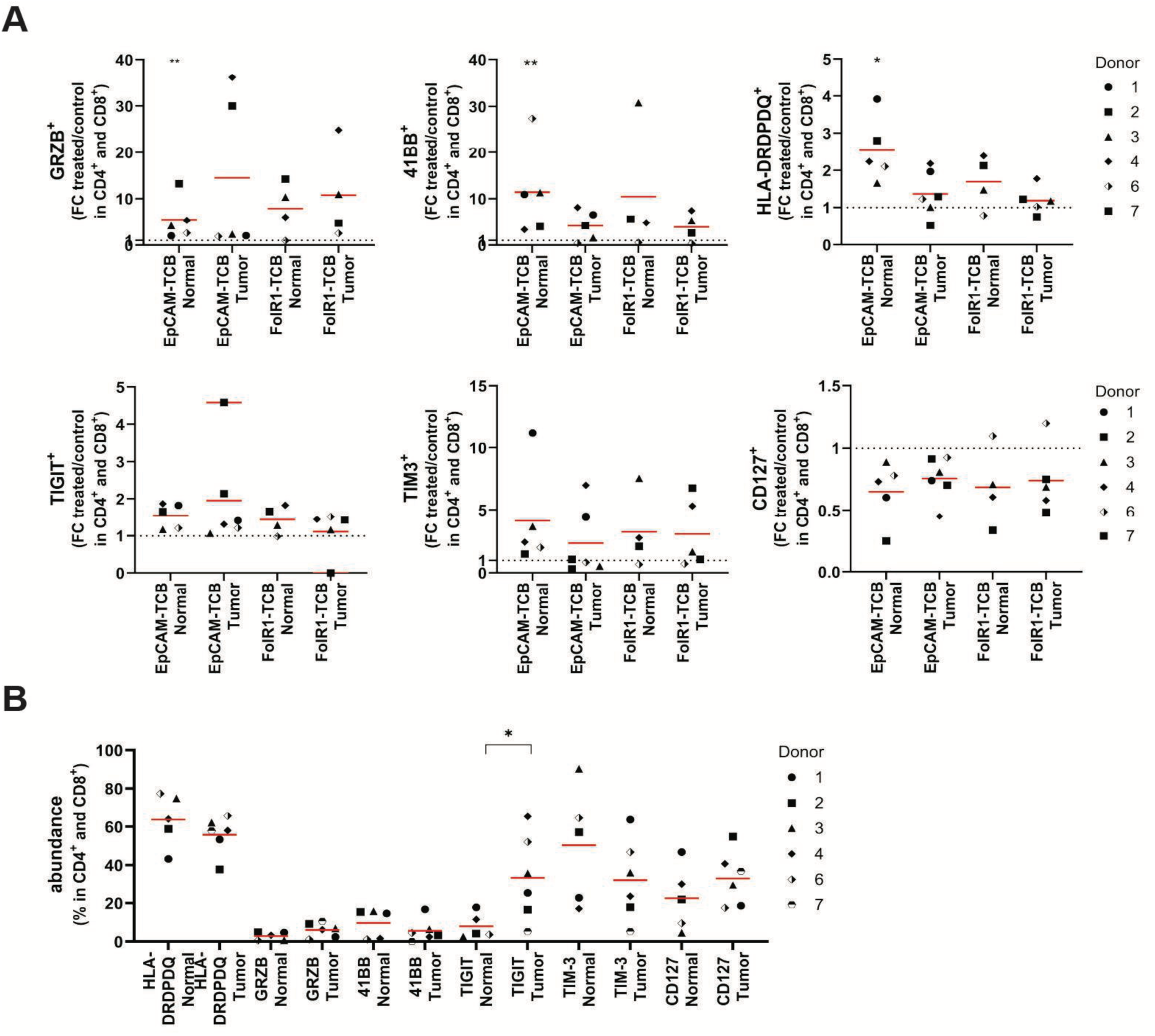
(A) Inter-donor variability in marker expression upon TCB treatments in CD4+ and CD8+ T cells normal and matched tumor explants for (top left to bottom right) GrzB, 4-1BB, HLA-DRDPDQ, CD127, TIM-3 and TIGIT. (B) Inter-donor variability of select markers in CD4+ and CD8+ T cells in lung baseline tissues. Red lines indicate the distribution mean.

**Supplementary Figure 3.**
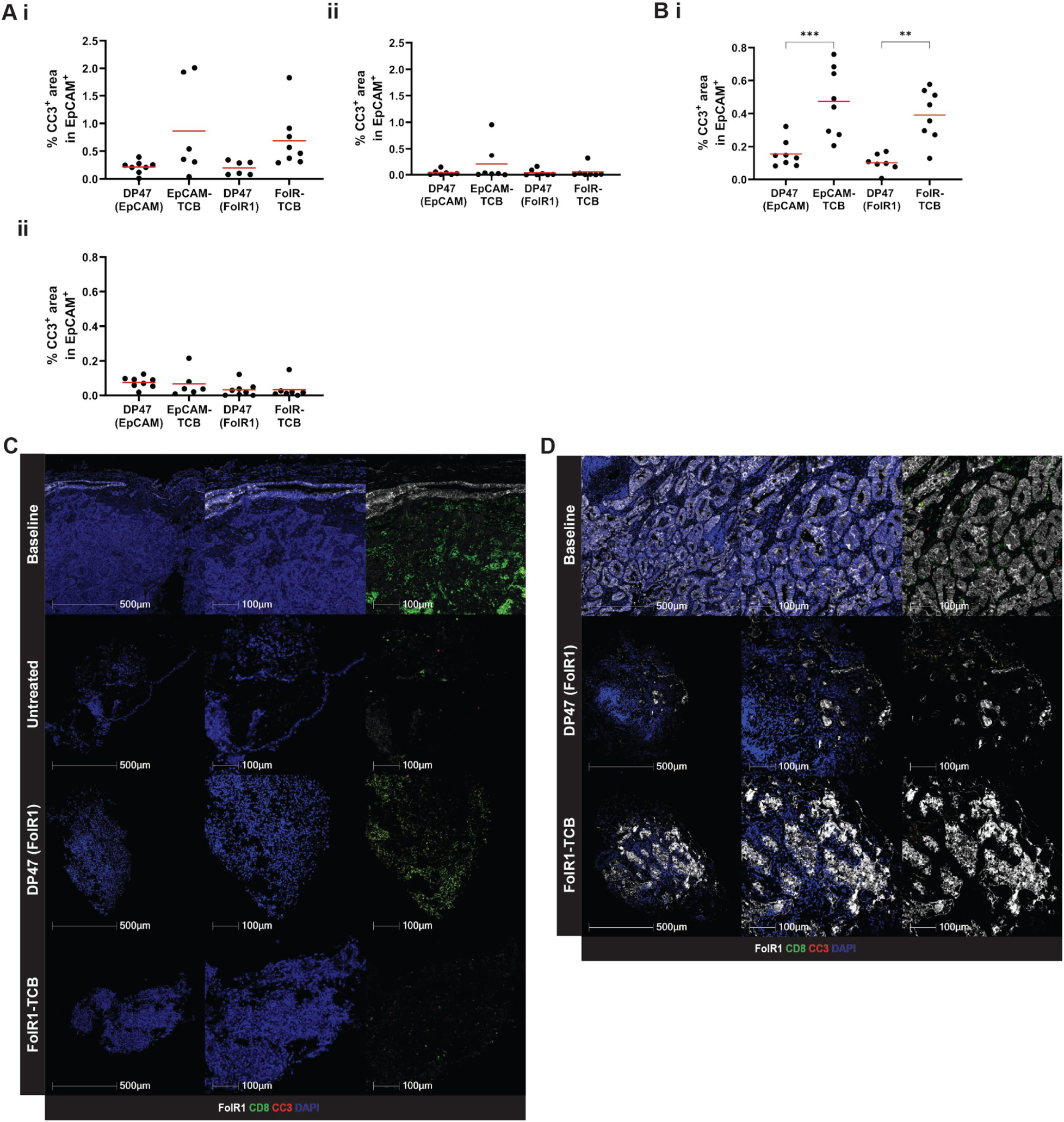
**CC3-positive areas detected in lung explants across donors**. Intra-donor variability of CC3-positive epithelial area in baseline, vehicle-treated and TCB-treated explants from donor 2 (A i, normal and A ii, tumor) and donor 4 (B i, normal and B ii, tumor) . Representative images of (C) donor 2 and (D) donor 4 tumor baseline tissue and explants showing FolR1-expressing cells, CD8 T cells and CC3 staining. Red lines indicate the distribution mean.

**SI Table 1.** Donor characteristics and explant culture features.

| Donor ID | Tissue | Source | Clinical Information | Treatment status | Immune infiltration into tumor sample | Tumor cells (% viable epithelial cells) | Necrosis/apoptosis (% whole tissue area) | Culture time ex vivo [hr] |
| --- | --- | --- | --- | --- | --- | --- | --- | --- |
| 1 | lung | Cytes Bio | 83 y/o, male, squamous carcinoma | naïve | excluded | 99 | 45 | 48 |
| 2 | lung | Cytes Bio | 66 y/o, male, typical carcinoid (neuroendocrine) tumor | naïve | deserted | 99 | 0 | 24 |
| 3 | lung | Cytes Bio | 45 y/o, male, lung nodule | naïve | excluded | 100 | 0 | 24 |
| 4 | lung | Humanitas | 74 y/o, male, adenocarcinoma | naïve | excluded | 100 | 2 | 48 |
| 6 | lung | HTCR | 70 y/o male lung nodule | naïve | could not be assessed | 10 | 60 | 24 |
| 7 | lung | HTCR | 78 y/o, female, typical carcinoid (neuroendocrine) tumor | naïve | deserted | 99 | 0 | 24 |
| 10 | colon | Humanitas | 51 y/o, male, adenocarcinoma | naïve |  |  |  | 48 |
| 11 | colon | DBM | 81 y/o, male, adenocarcinoma | naïve |  |  |  | 24 |
| 12 | colon | Humanitas | 73 y/o, female, adenocarcinoma | naïve |  |  |  | 24 |
| 13 | colon | HTCR | 45 y/o, female, adenocarcinoma | naïve |  |  |  | 24,48 |

**SI Table 2.** Lymphocyte and tumor cell markers used in flow cytometry experiments.

| Marker | Vendor, Cat number | Clone | Panel |
| --- | --- | --- | --- |
| CD45 | BD Biosciences, 564106 | HI30 | lymphocyte |
| CD3 | BD Biosciences, 612896 | UCHT1 | lymphocyte |
| CD4 | BD Biosciences, 612748 | SK3 | lymphocyte |
| TIM-3 | BD Biosciences, 752363 | 7D3 | lymphocyte |
| HLA-DRDPDQ | BD Biosciences, 741157 | Tu39 | lymphocyte |
| Ki67 | BD Biosciences, 564071 | B56 | lymphocyte |
| CD69 | BD Biosciences, 748764 | FN50 | lymphocyte |
| CD56 | BD Biosciences, 564058 | NCAM16.2 | lymphocyte |
| CD25 | Biolegend, 302640 | BC96 | lymphocyte |
| CD8 | Biolegend, 301044 | RPA-T8 | lymphocyte |
| CD45RA | Biolegend, 304134 | HI100 | lymphocyte |
| 41BB | Biolegend, 309820 | 4B4-1 | lymphocyte |
| FoxP3 | Biolegend, 320212 | 259D | lymphocyte |
| PD-1 | Biolegend, 329918 | EH12.2H7 | lymphocyte |
| CD127 | Biolegend, 351336 | A019D5 | lymphocyte |
| Granzyme B | Biolegend, 372222 | QA16A02 | lymphocyte |
| CD39 | Biolegend, 328210 | A1 | lymphocyte |
| Zombie NIR | Biolegend, 423105 | NA | lymphocyte |
| TIGIT | Thermo Fisher Scientific, 12-9500-42 | MBSA43 | lymphocyte |
| PD-L1 | BD Biosciences, 741881 | MIH1 | tumor |
| CD86 | BD Biosciences, 748377 | BU63 | tumor |
| CD11C | BD Biosciences, 741139 | B-ly6 | tumor |
| CD19 | BD Biosciences, 562947 | SJ25C1 | tumor |
| FoIR | BD Biosciences, 569812 | Clone 5 | tumor |
| CD11B | BD Biosciences, 564454 | M1/70 | tumor |
| CC3 | BD Biosciences, 570179 | C92-605.rMAb | tumor |
| CD45 | BD Biosciences, 563792 | HI30 | tumor |
| CD3 | Biolegend, 317330 | OKT3 | tumor |
| CD68 | Biolegend, 333814 | Y1/82A | tumor |
| CD14 | Biolegend, 325618 | HCD14 | tumor |
| CD90 | Biolegend, 328134 | 5E10 | tumor |
| EpCAM | Biolegend, 324206 | 9C4 | tumor |
| Zombie UV | Biolegend, 423107 | NA | tumor |
| CD31 | Bio-Rad, MCA1738SBUV665 | WM59 | tumor |
| YD TCR | Miltenyi Biotec, 130-111-011 | REA771 | tumor |
| CEACAM5 | in-house; labelled with AF750 | - | tumor |

## Methods

### Tissue collection

Tumor and adjacent normal tissue resections from lung and colon cancer patients were obtained from four of sources: 1. HTCR tissues: Human tissue samples and annotated clinical data were obtained from patients undergoing tumor resections within the framework of the nonprofit foundation Human Tissues and Cell Research (HTCR) (Reference: doi: 10.1023/A:1026392429112). The framework of the HTCR Foundation, which includes written informed consent from all donors, has been reviewed by the ethics commission of the Faculty of Medicine, Ludwig Maximilians University (no. 025-12) and the Bavarian State Medical Association (no. 11142). 2. DMB tissues: Human tissues and associated clinical information were obtained from patients undergoing tumor resection at the University Center for Gastrointestinal and Liver Disease (Clarunis), Basel, Switzerland. Tissues were sourced through a collaborative framework adhering to legal and ethical regulations. The project was performed in accordance with the Helsinki Declaration and reviewed and approved by the Ethics Committee of Basel, EKBB, no. 2019-02118. 3. Humanitas tissues: tissues and associated clinical information were obtained from patients undergoing tumor resection at the Humanitas Research Hospital, Milan, Italy. Samples were sourced through a collaborative framework adhering to legal and ethical regulations. The project was reviewed and approved by Comitato Etico Terrirotiale Lombardia 5 (Ethical approval 3631). 4. BeCytes Biotechnologies tissues: Lung samples were collected as part of the project “Modelling cancer-immunotherapy–induced pulmonary toxicities with in vitro human assays”, implemented at MútuaTerrassa University Hospital (code P/22-002/MCIPT.2022.v1). The study was approved by the Research Ethics Committee on Medicinal Products in ordinary session no. 02/2022 on February 23, 2022, and by the Research Ethics Committee on Medicinal Products (CEIm) of Parc Taulí de Sabadell (Barcelona, Spain) in ordinary session no. 2022/5030 on March 29, 2022. Additional lung samples were collected at Josep Trueta University Hospital of Girona (code 2023.19), following approval by its Research Ethics Committee on Medicinal Products in ordinary session no. 13/2023 on October 9, 2023. Samples were received on ice in transport medium within 24 h from surgery and processed immediately.

### Tissue processing and explant culture

Tissue was processed as described in ^20,43^, with minor modifications. Briefly, lung tissues were washed in wash buffer (Hank’s balanced salt solution (Gibco, 24020-091) and 0.2 mg/ml Primocin (Invivogen, ant-pm-1), to remove excess blood and placed in a glass petri dish (VWR, 391-0581) on ice and manually cut with a surgical scalpel into small fragments of about 1 mm^3^ (explants). For colon tissues, resections were washed and the mucosal layer separated from the underlying submucosa and muscle tissue, which were discarded. Mucus was removed from the mucosal layer by scraping the epithelial surface gently with the dull side of a scalpel blade, after which tissues were washed in wash buffer again and cut into explants. In the meantime, a 96-well plate (Greiner, 655098) was pre-coated with 30 μl of a collagen-Matrigel mixture (to a final concentration of 0.75 % sodium bicarbonate (Gibco, 25080-094), 1 mg/ml rat tail Collagen I High Concentration (Sigma-Aldrich, 354249), 2 mg/ml Matrigel® Matrix High Concentration (Corning, 354262) in tumor medium). Tumor medium comprised DMEM high glucose (Gibco, 11960) supplemented with 1 mM of sodium pyruvate (Gibco, 11360-070), 0.1 mM MEM non-essential amino acids (NEAA) (Gibco, 11140-050), 2 mM of GlutaMAX (Gibco, 35050-038), 10% heat-inactivated FBS (Sigma-Aldrich, F4135), 100 U/ml penicillin and 100 μg/ml streptomycin (Gibco, 15140-122). The matrix mixture was gelled in the incubator at 37°C for 30 minutes, after which explants were distributed randomly (1 explant per well) using tweezers and topped with an additional 20 μl of collagen-Matrigel mixture per well. After gelation of this second matrix volume, 150 μl of medium, with or without treatment, were added per well. Explants were incubated at 37°C, 5% CO2 and 90% humidity (see SI Table 1 for culture times).

### TCB ex vivo treatment

Targeting T cell TCBs included FolR1 TCB, CEACAM5 TCB and EpCAM TCB; non-targeting (vehicle) DP47 controls included a control for FolR1 TCB and another control for EpCAM TCB; the latter also served as vehicle control for experiments with CEACAM5 TCB, given the analogous protein structure. All TCBs were generated by Hoffman-La Roche as described elsewhere ^8,44^. TCB concentrations used across experiments were 0.5 μg/ml except where stated otherwise.

### Tissue dissociation

Fresh, uncultured “baseline” tissue was finely chopped using a scalpel, while cultured explants w*ere harv*ested from the 96-well plate by removing them from the collagen-Matrigel matrix with forceps and pooling a minimum of 8 fragments per treatment condition. The chopped baseline tissue and the explant pools were transferred into separate wells of a 24-well plate (TPP, 92024) containing digestion medium (50 % MACS Tissue Storage Solution (Miltenyi Biotec, 130-100-008), 45 % Accutase (Sigma-Aldrich, A6964), 33% bovine serum albumin (Sigma-Aldrich, H6254), 275 U/ml collagenase IV (Gibco, 17104-019), 10 U/ml DNase I Type 4 (Roche, 04536282001) and 471 U/ml hyaluronidase (Sigma-Aldrich, H6254)). Samples were digested in this medium on a shaker (Benchmark Scientific, BT1502) at 300-350 rpm for 30 minutes at 37°C. The resulting single cell suspension was filtered through a 70 μM strainer (Miltenyi Biotec, 130-098-462) and the yield was maximised by gently mashing remaining tissue explants against the strainer with a syringe plunger (BD Biosciences, 300912). Samples were washed once by adding RPMI 1640 (Gibco, 31870-025) to the filtered cell suspension, followed by a centrifugation step for 5 minutes at 400 g. If substantial amounts of red blood cells were present within the resulting pellets, a lysis step was performed by resuspending the pellet in Red Blood Cell (RBC) Lysis Buffer (Roche, 11814389001) for 15 minutes at 4°C, followed by an additional washing step as described above. Digest from the baseline tissue was then cryopreserved in pZerve (Sigma-Aldrich, Z1653) and stored in liquid nitrogen until processing for flow cytometry. Digests from explants after treatment were directly stained and processed for flow cytometry.

### Flow Cytometry calibrator sample

In order to generate a reference sample (calibrator sample) to be used across flow cytometry experiments, the following cellular fractions were pooled: (1) The human gastric adenocarcinoma MKN45 (DSMZ, ACC 409) and the breast cancer T47D (ATCC, HTB-133) cell lines were cultured at 37°C, 5% CO2 and 90% humidity in RPMI1640 medium (Thermo Fisher Scientific, 11875093) supplemented with 10% FBS, 2 mM Glutamax, 0.1 mM NEAA, 100 U/ml penicillin, 100 μg/ml streptomycin and 1 mM sodium pyruvate. Both cell lines were harvested before confluency was reached, combined in a 1:1 ratio and cryopreserved in the culture medium, which was further supplemented with 10% FBS and 10% DMSO (Thermo Fisher Scientific, J66650.AK), at a density of 6 x 10^5^ cells per vial. (2) PBMCs were isolated from a buffy coat (Blutspende Zürich) using Lymphoprep solution (StemCell, 07801) according to the manufacturer’s protocol. Half of the obtained PBMCs were stimulated according to Thermo Fisher’s T cell activation protocol using anti-CD3 and anti-CD28 antibodies (Thermo Fisher Scientific). Briefly, a 6-well plate (TPP, 92006) was coated with 2ml of PBS containing 1 μg/ml anti-CD3 antibody (Biolegend, Cat. 16-0039-85) for 2-3 h at 37°C. The PBS was then removed and without washing, 2ml of RPMI1640 medium (supplemented with 2 mM Glutamax, 100 U/ml penicillin, 100 μg/ml streptomycin and 10% FBS) containing 3 x 10^6^ cells/ml PBMCs and 3 μg/ml anti-CD28 (Biolegend, 302923) was added to each well. Plates were cultured for 24h at 37°C and 5% CO_2_. Following incubation, stimulated and unstimulated PBMCs were harvested, combined in a 1:1 ratio and cryopreserved in 90% FBS and 10% DMSO at a density of 1.2 x 10^6^ cells/vial. On the day of staining, one vial of MKN45/T47D cells and one vial of PBMCs were thawed and pooled. Final calibrator samples contained 1.8 million cells in a 2:2:1:1 ratio of stimulated PBMCs, unstimulated PBMCs, MKN45 cells and T47D cells.

### Flow cytometry

Digests from baseline (thawed) and explants (fresh), as well as calibrator samples, were analyzed for lymphocyte and tumor markers (SI Table 2). In short, digests were washed twice in PBS (Gibco, 10010-023) and transferred to a 96-well V-bottom plate (Corning, 3894), with all centrifugation steps performed at 400 x g and 4°C for 5 minutes, and all staining steps conducted at room temperature in the dark. Viability dyes Zombie UV or Zombie NIR were diluted 1:400 in PBS and cells were stained with 50 μl/well of the viability dye for 20 min. After one washing step with 100 μl/well FACS buffer (demineralized water plus 1x PBS (VWR, 392-0440), 2% FBS (Sigma, F4135), 5 mM EDTA (VWR, E177), 7.5 mM NaN3 (Sigma, S2002)), cells were incubated for 10 min with 25 μl/well Fcx block (Biolegend, 422301) diluted 1:20 in FACS buffer. Without any washing step, 25 μl/well of a 2x staining mix for extracellular markers (1:50 in FACS buffer) was added and the cells were incubated for 20 minutes. After a further washing step using FACS buffer, cells were fixed and permeabilized using 50 μl/well of the eBioscience™ Foxp3 / Transcription Factor Staining Buffer Set (Thermo Fisher Scientific, 00-5523-00), according to the manufacturer’s protocol, for 45 minutes at room temperature or overnight at 4°C. Cells were then washed with permeabilization/wash solution (Thermo Fisher Scientific, 00-5523-00) and stained with 50 μl of a staining mix for intracellular/intranuclear markers (1:100) for 2 hours at room temperature or overnight at 4°C, followed by an additional wash step in permeabilization/wash solution. To correct for fluorescence spillover, single-stain compensation controls were prepared by individually staining compensation beads (Invitrogen, 01-3333-42) with each antibody, while ArC Amine Reactive Compensation Beads (Invitrogen, A10346) were used for the Zombie UV and Zombie NIR viability dyes. Flow cytometry data was acquired on a Symphony A3 (BD Biosciences) and data analysis was performed with FlowJo (version 10.10.0).

### Cytokine and chemokine analysis

Explant culture supernatants were collected at endpoint after treatment with EpCAM TCB (donors 1-4, 7), FolR1 TCB (donors 2-4, 7), or CEACAM5 TCB (donors 8-11), along with their respective DP47 non-targeting controls. For each condition, supernatants from a minimum of 8 explants were pooled and stored at −80°C until analysis, which was carried out using LEGENDplex™ multiplex bead-based immunoassay kits (Biolegend), including the Human CD8/NK Panel V02 (741187), the Human Proinflammatory Chemokine Panel 1 (740985) and the Human T Helper Cytokine Panels Version 2 (741028). All procedures followed the manufacturer’s protocol, with the modification of using half the specified volume for all reagents and samples. Data was acquired on a Symphony A3 flow cytometer (BD Biosciences) and analyzed using Biolegend’s Data Analysis Software Suite for LEGENDplex™.

### FFPE Embedding and multiplexed immunofluorescence

Tissues designated for immunohistochemistry (IHC) and multiplexed immunofluorescence (mIF) were fixed in 4% paraformaldehyde (PFA; Sigma, HT501128) at 4°C overnight and subsequently stored in PBS prior to paraffin embedding. Histological processing was facilitated using an in-house 3D printed 48-well Histogel mold as described previously ^45^, enabling alignment of multiple tissue fragments within a single formalin-fixed paraffin-embedded (FFPE) block. The histoarray was solidified at 4°C for 10 minutes, extracted from the mold, and fixed explants were positioned within each well using tweezers and supplemented with additional Histogel. The completed histoarray was then transferred to a labeled histology cassette (Rotilabo). Tissue processing was carried out using the Tissue-Tek VIP 6 AI system (Sakura), using xylene (Sigma, 24764-2) and ethanol (Fluka, 02854-2.5L), while paraffin embedding was performed using the Leica HistoCore Arcadia H system with paraffin (Leica, X880.2). FFPE blocks were sectioned to a thickness of 3 µm using a Leica RM2235 microtome (Leica Biosystems). H&E staining was performed with the HistoCore SPECTRA ST stainer (Leica Biosystems). mIF staining was performed using a Ventana Discovery Ultra automated tissue stainer (Roche Tissue Diagnostics). Slides were baked at 60°C for 8 minutes, followed by two deparaffinization steps at 69°C for 8 minutes each. Heat-induced antigen retrieval was carried out at 92°C for 32 minutes using Tris-EDTA buffer (pH 7.8; Ventana, 950-500). Blocking steps, each lasting 16 minutes, employed Discovery Inhibitor (Ventana, 760-4840), followed by neutralization post-blocking. Primary antibody detection utilized horseradish peroxidase-conjugated secondary antibodies (OmniMap anti-rabbit HRP, Ventana, 760-4311). Corresponding Opal dyes (Akoya Biosciences) were applied sequentially for marker visualization. After each cycle of primary antibody, secondary antibody, and Opal dye application, slides underwent antibody neutralization and horseradish peroxidase (HRP) denaturation before initiating subsequent cycles, using Discovery Inhibitor. Specific markers stained were: CD8 (ab4055, Abcam; dilution 1:200), Cleaved Caspase 3 (9661S, Cell Signaling Technology; dilution 1:100), CD45 (13917S, Cell Signaling Technology; dilution 1:50), EpCAM (ab213500, Abcam; dilution 1:100), FOLR1 (NBP2-61773, Novus Biologicals; dilution 1:1000), and CEACAM5 (ab207718, Abcam; dilution 1:500). Finally, slides were counterstained with 4′,6-diamidino-2-phenylindole (DAPI). Slides were coverslipped using a CTM6 Coverslipper (Histocom AG) and mounted with Immu-Mount (Epredia, 9990414).

### Scanning and quantification

IHC slides were scanned at 20x magnification using the Olympus SLIDEVIEW VS200 slide scanner. Multiplex immunofluorescence slides were scanned using the PhenoImager HT system (Akoya Biosciences). Quantification was performed using Halo AI software (version 3.6) with user-guided artificial intelligence-based tissue segmentation. Specifically, mIF images were manually annotated and tissue areas assigned into three distinct categories: “background,” “tissue,” and “epithelium.” The “background” category included all dark areas devoid of tissue (e.g. surrounding explants or within alveolar spaces). EpCAM-positive tissue regions exhibiting typical, non-necrotic morphology were classified as “epithelium”, while “tissue” comprised all non-epithelium, non-background areas. Based on these annotations in at least three baseline and three explant images, independent classification models (DensNet V2 classifier, Halo AI) were trained separately for lung and colon samples. This step ensured that the algorithm could appropriately account for inter- and intra-patient heterogeneity. Within the segmented regions, the positively stained areas for different markers were quantified. For CC3, EpCAM and CEACAM5 a user-defined threshold was applied to exclude background and nonspecific signal. For calculating tissue cellularity in mIF images, a minimum of four tissue areas were defined in DP47-treated explants using manual annotation or in baseline tissues, using circular ROIs of standard size. Within these areas, the area occupied by DAPI signal was calculated using a standard HALO Algorithm (Indica Labs - Area Quantification FL.v2.3.4), based on manually adjusted DAPI signal thresholds.

### Immune Infiltration status of baseline tissues

Tumor tissues from donors 1-7 were classified as infiltrated, excluded or deserted by a human pathologist using a published method ^46^, which was however applied to H&E sections. Additionally, the percentage of viable tumor cells and the percentage of necrosis/apoptosis levels within tumors were also assessed.

### Statistical analysis and graph generation

All statistical analyses were performed using GraphPad Prism (Dotmatics, v10.3.0) and R Statistical Software (Core Team 2023, v4.3.3). Data are presented as mean ± standard deviation unless otherwise specified. All analyses include data from a minimum of n=3 tissues except for SI Fig 1A, where explant cohorts from n=2 normal tissues were included. Outliers in flow cytometry datasets were identified and removed in GraphPad Prism using the ROUT method (Q=1%). Data were tested for normality using the Shapiro-Wilk test; significance in normally distributed data was then assessed using an unpaired, two-tailed t-test (for 2-sample comparisons) or a one-way ANOVA followed by a Tukey’s pos-hoc test (for comparisons of 3 samples or more), while significance in data lacking a normal distribution was assessed using a Mann-Whitney U test (2 samples) or a Kruskal-Wallis Test followed by a Dunn’s post-hoc test (3 samples or more). P-values < 0.05 were considered statistically significant (*p < 0.05, **p < 0.01, ***p < 0.001, ****p < 0.0001). Experimental workflows depicted in Fig. 1A [publication licence] and Fig. 2A [publication licence] were generated with BioRender (BioRender.com). All analyses were carried out for every donor, unless stated otherwise in the figure legend.

## Acknowledgements

We acknowledge the support of the nonprofit foundation HTCR, which holds human tissue on trust, making it broadly available for research on an ethical and legal basis. We extend our gratitude to BeCytes Biotechnologies, a BioIVT Company and Dr. Mireia Serra Mitjants from the Thoracic Surgery Service at MútuaTerrassa University Hospital (Terrassa, Spain); the Donation and Transplantation Institute (DTI, Spain); and Dr. Matilde Rubio and Dr. Xavier Baldo from the Thoracic Surgery Service at Josep Trueta University Hospital (Girona, Spain) for facilitating access to biospecimens and ensuring their availability for research in compliance with ethical and legal standards, following the provision of informed consent by the donors. We thank Salvatore Piscuoglio and his team at the University Hospital Basel and Humanitas Research Hospital for providing the tissues and clinical data. We would further like to express our gratitude to Dr. Sylvia Herter for her generous and valuable input and feedback.

## Author contributions

Study idea and design by E.D., M.L and L.C. Manuscript writing by E.D., M.T., T.J., C.Y., N.N., L.C. and M.L. Data generation by M.T. and T.J. HALO AI quantification by M.T. Flow Cytometry quantification by T.J and C.Y. Cellularity analysis in mIF images by E.D. Optimization of mIF histology stainings by M.O. Histological assessment by A.B.

## References

1. Anderson, G. S. F. & Chapman, M. A. T cell-redirecting therapies in hematological malignancies: Current developments and novel strategies for improved targeting. Mol. Ther. 32, 2856–2891 (2024).

2. Albayrak, G., Wan, P. K.-T., Fisher, K. & Seymour, L. W. T cell engagers: expanding horizons in oncology and beyond. Br. J. Cancer 1–9 (2025) doi:10.1038/s41416-025-03125-y.

3. Tapia-Galisteo, A., Álvarez-Vallina, L. & Sanz, L. Bi- and trispecific immune cell engagers for immunotherapy of hematological malignancies. J. Hematol. Oncol. 16, 83 (2023).

4. Liu, J. & Zhu, J. Progresses of T-cell-engaging bispecific antibodies in treatment of solid tumors. Int. Immunopharmacol. 138, 112609 (2024).

5. O’Brien, E., Mayer, C. M. & Arend, R. C. Exploring T-cell bispecific antibodies in gynecologic malignancy. Gynecol. Oncol. Rep. 59, 101772 (2025).

6. Edeline, J., Houot, R., Marabelle, A. & Alcantara, M. CAR-T cells and BiTEs in solid tumors: challenges and perspectives. J. Hematol. Oncol. 14, 65 (2021).

7. Kebenko, M. et al. A multicenter phase 1 study of solitomab (MT110, AMG 110), a bispecific EpCAM/CD3 T-cell engager (BiTE®) antibody construct, in patients with refractory solid tumors. OncoImmunology 7, e1450710 (2018).

8. Kerns, S. J. et al. Human immunocompetent Organ-on-Chip platforms allow safety profiling of tumor-targeted T-cell bispecific antibodies. eLife 10, e67106 (2021).

9. Powley, I. R. et al. Patient-derived explants (PDEs) as a powerful preclinical platform for anti-cancer drug and biomarker discovery. Br. J. Cancer 122, 735–744 (2020).

10. Genta, S., Coburn, B., Cescon, D. W. & Spreafico, A. Patient-derived cancer models: Valuable platforms for anticancer drug testing. Front. Oncol. 12, 976065 (2022).

11. Meijer, T. G., Naipal, K. A., Jager, A. & Gent, D. C. van. Ex vivo tumor culture systems for functional drug testing and therapy response prediction. Futur. Sci. OA 3, FSO190 (2017).

12. Junk, D. et al. Human tissue cultures of lung cancer predict patient susceptibility to immune-checkpoint inhibition. Cell Death Discov. 7, 264 (2021).

13. Turpin, R. et al. Patient-derived tumor explant models of tumor immune microenvironment reveal distinct and reproducible immunotherapy responses. OncoImmunology 14, 2466305 (2025).

14. Sharon, S. et al. Explant Modeling of the Immune Environment of Head and Neck Cancer. Front. Oncol. 11, 611365 (2021).

15. Templeton, A. R. et al. Patient-Derived Explants as a Precision Medicine Patient-Proximal Testing Platform Informing Cancer Management. Front. Oncol. 11, 767697 (2021).

16. Naipal, K. A. T. et al. Tumor slice culture system to assess drug response of primary breast cancer. BMC Cancer 16, 78 (2016).

17. Gerlach, M. M. et al. Slice cultures from head and neck squamous cell carcinoma: a novel test system for drug susceptibility and mechanisms of resistance. Br. J. Cancer 110, 479–488 (2014).

18. Demetriou, C. et al. An optimised patient-derived explant platform for breast cancer reflects clinical responses to chemotherapy and antibody-directed therapy. Sci. Rep. 14, 12833 (2024).

19. Gavert, N. et al. Ex vivo organotypic cultures for synergistic therapy prioritization identify patient-specific responses to combined MEK and Src inhibition in colorectal cancer. *Nat*. Cancer 3, 219–231 (2022).

20. Voabil, P. et al. An ex vivo tumor fragment platform to dissect response to PD-1 blockade in cancer. Nat. Med. 27, 1250–1261 (2021).

21. Shekarian, T. et al. Immunotherapy of glioblastoma explants induces interferon-γ responses and spatial immune cell rearrangements in tumor center, but not periphery. Sci. Adv. 8, eabn9440 (2022).

22. Fernando, K. et al. Extended human lymph node explants for evaluation of adaptive immunity. Trends Biotechnol. (2025) doi:10.1016/j.tibtech.2025.07.020.

23. Kamer, I. et al. Immunotherapy response modeling by ex-vivo organ culture for lung cancer. *Cancer Immunol.*, Immunother. 70, 2223–2234 (2021).

24. Kaptein, P. et al. Reinvigoration of translational activity in dysfunctional T cells initiates the early intratumoral response to PD-1 blockade. bioRxiv 2025.09.24.676875 (2025) doi:10.1101/2025.09.24.676875.

25. Autrup, H. et al. Explant culture of human colon. Gastroenterology 74, 1248–1257 (1978).

26. Lee-Ferris, R. E. et al. Prolonged airway explant culture enables study of health, disease, and viral pathogenesis. Sci. Adv. 11, eadp0451 (2025).

27. Kauer, J. et al. CD18 Antibody Application Blocks Unwanted Off-Target T Cell Activation Caused by Bispecific Antibodies. Cancers 13, 4596 (2021).

28. Kiniry, B. E. et al. Differential Expression of CD8+ T Cell Cytotoxic Effector Molecules in Blood and Gastrointestinal Mucosa in HIV-1 Infection. J. Immunol. 200, 1876–1888 (2018).

29. Thompson, R. & Cao, X. Reassessing granzyme B: unveiling perforin-independent versatility in immune responses and therapeutic potentials. Front. Immunol. 15, 1392535 (2024).

30. Ogata, M. et al. Granzyme B-dependent and perforin-independent DNA fragmentation in intestinal epithelial cells induced by anti-CD3 mAb-activated intra-epithelial lymphocytes. Cell Tissue Res. 352, 287–300 (2013).

31. Motyka, B. et al. Mannose 6-Phosphate/Insulin-like Growth Factor II Receptor Is a Death Receptor for Granzyme B during Cytotoxic T Cell–Induced Apoptosis. Cell 103, 491–500 (2000).

32. Kurschus, F. C. et al. Killing of target cells by redirected granzyme B in the absence of perforin. FEBS Lett. 562, 87–92 (2004).

33. Tang, J. et al. B Cells and Tertiary Lymphoid Structures Influence Survival in Lung Cancer Patients with Resectable Tumors. Cancers 12, 2644 (2020).

34. Shu, L., Tao, T., Xiao, D., Liu, S. & Tao, Y. The role of B cell immunity in lung adenocarcinoma. Genes Immun. 26, 253–265 (2025).

35. Schacht, S.-S. et al. Activation and maturation of antigen-specific B cells in nonectopic lung infiltrates are independent of germinal center reactions in the draining lymph node. Cell. Mol. Immunol. 22, 612–627 (2025).

36. Cao, L., Leclercq-Cohen, G., Klein, C., Sorrentino, A. & Bacac, M. Mechanistic insights into resistance mechanisms to T cell engagers. Front. Immunol. 16, 1583044 (2025).

37. Hickey, J. W. et al. T cell-mediated curation and restructuring of tumor tissue coordinates an effective immune response. Cell Rep. 42, 113494 (2023).

38. Maggi, E. et al. T cell landscape in the microenvironment of human solid tumors. Immunol. Lett. 270, 106942 (2024).

39. Schenkel, J. M. & Pauken, K. E. Localization, tissue biology and T cell state — implications for cancer immunotherapy. Nat. Rev. Immunol. 23, 807–823 (2023).

40. Mowat, A. McI. Anatomical basis of tolerance and immunity to intestinal antigens. Nat. Rev. Immunol. 3, 331–341 (2003).

41. Hegde, P. S. & Chen, D. S. Top 10 Challenges in Cancer Immunotherapy. Immunity 52, 17–35 (2020).

42. Abdelmotaleb, O., Schneider, A., Gassner, C., Märsch, S. & Klein, C. The impact of CD3 affinity-attenuation on T cell engaging bispecific antibodies: is it really that simple? Expert Opin. Drug Discov. 20, 943–949 (2025).

43. Roelofsen, L. M. et al. Protocol for ex vivo culture of patient-derived tumor fragments. STAR Protoc. 4, 102282 (2023).

44. Harter, M. F. et al. Analysis of off-tumour toxicities of T-cell-engaging bispecific antibodies via donor-matched intestinal organoids and tumouroids. *Nat*. Biomed. Eng. 8, 345–360 (2024).

45. Harter, M. F. et al. High-throughput histopathology for complex in vitro models. bioRxiv 2025.04.01.646522 (2025) doi:10.1101/2025.04.01.646522.

46. Li, X. et al. Automated tumor immunophenotyping predicts clinical benefit from anti-PD-L1 immunotherapy. J. Pathol. 263, 190–202 (2024).

